# Recovery of Metagenomic Data from the *Aedes aegypti* Microbiome using a Reproducible Snakemake Pipeline: MINUUR

**DOI:** 10.1101/2022.08.09.503283

**Authors:** Aidan Foo, Louise Cerdeira, Grant L. Hughes, Eva Heinz

## Abstract

**Background:** Ongoing research of the mosquito microbiome aims to uncover novel strategies to reduce pathogen transmission. Sequencing costs, especially for metagenomics, are however still significant. A resource that is increasingly used to gain insights into host-associated microbiomes is the large amount of publicly available genomic data based on whole organisms like mosquitoes, which includes sequencing reads of the host-associated microbes and provides the opportunity to gain additional value of these initially host-focused sequencing projects.

**Methods:** To analyse non-host reads from existing genomic data, we developed a snakemake workflow called MINUUR (**M**icrobial **IN**sights **U**sing **U**nmapped **R**eads). Within MINUUR, reads derived from the host-associated microbiome were extracted and characterised using taxonomic classifications and metagenome assembly followed by binning and quality assessment. We applied this pipeline to five publicly available *Aedes aegypti* genomic datasets, consisting of 62 samples with a broad range of sequencing depths.

**Results:** We demonstrate that MINUUR recovers previously identified phyla and genera and is able to extract bacterial metagenome assembled genomes (MAGs) associated to the microbiome. Of these MAGS, 42 are high-quality representatives with >90% completeness and <5% contamination. These MAGs improve the genomic representation of the mosquito microbiome and can be used to facilitate genomic investigation of key genes of interest. Furthermore, we show that samples with a high number of KRAKEN2 assigned reads produce more MAGs.

**Conclusions:** Our metagenomics workflow, MINUUR, was applied to a range of *Aedes aegypti* genomic samples to characterise microbiome-associated reads. We confirm the presence of key mosquito-associated symbionts that have previously been identified in other studies and recovered high-quality bacterial MAGs. In addition, MINUUR and its associated documentation are freely available on GitHub (https://github.com/aidanfoo96/MINUUR) and provide researchers with a convenient workflow to investigate microbiome data included in the sequencing data for any applicable host genome of interest.

## Introduction

*Aedes aegypti* is an important vector of human pathogens including dengue virus (DENV), yellow fever virus, chikungunya virus and Zika virus. DENV cases alone are estimated to cause 10,000 deaths and 100 million infections per year, contributing to significant burden of human morbidity and mortality worldwide (1). Interest in the mosquito microbiome has emerged due to evidence of its influence in vectorial capacity (2,3), offering potential for novel approaches to reduce pathogen transmission from mosquitoes to vertebrate hosts (2,4–7).

Mosquito microbiomes are highly variable, dependent on multiple deterministic processes such as the environment (8–12), season (13), host factors (14,15), microbial interactions (16–18) and mosquito-microbe interactions (19–23). These findings from mosquito microbiome studies are largely driven by amplicon based 16S rRNA sequencing approaches (18,24,25). Metagenomic approaches for mosquito microbiome characterisation are limited in number (26,27), but would facilitate our understanding by providing the genomic context of symbionts (28). An attractive resource for gaining additional insights into microbiomes is to make use of the microbiome reads derived from whole genome sequencing (WGS), especially in cases where preparation of the host for sequencing included its associated microbiome. Studies in *Drosophila*, bumble bees, moths and nematodes have shown existing WGS data is a rich source to characterize associated symbionts (29–34). Whilst some studies include specific enrichment of non-host with bait sequences targeting a specific taxon of interest (29,32); it is also possible to assess microbes present in the sequencing experiment without prior enrichment, taking into account that in the latter case the microbiome recovered is likely a biased representation and lack of presence does not prove absence.

Whole genome shotgun sequencing is commonly used to study mosquito genomics at the individual (35,36) and population (37,38) level; meaning non-mosquito sequence data (we refer to these as non-reference reads, since they do not map to the reference genome of interest) are a source to identify mosquito microbiome members using metagenomics. Genomic surveillance programs such as the *Anopheles gambiae* 1000 Genomes Project contain a large number of genomic samples with each release (39) and, at time of writing, currently 100,514 *Aedes aegypti* whole-genome sequencing runs are deposited on the European Nucleotide Archive. As such, there is great potential to leverage existing mosquito WGS data to explore members of their microbiomes from non-reference reads.

To make use of this resource and streamline future large-scale analysis of mosquito metagenomes, we developed a Snakemake pipeline to provide **M**icrobial **IN**sight **U**sing **U**nmapped **R**eads from WGS data. MINUUR uses short read, whole genome sequencing data as input and performs an analysis of non-reference reads derived from a host sequencing experiment. We used MINUUR to study five published *Aedes aegypti* sequencing projects (in total 124 FASTQ files) currently available on the European Nucleotide Archive. We gained insight of associated microbes based on taxonomic read classifications and recover high-quality metagenome assembled genomes (MAGs) of mosquito-associated bacteria. To assess the suitability of samples for MAG recovery, we also investigated patterns between KRAKEN2 assigned reads and the number of MAGs within samples. We show samples with taxa assigned high numbers of KRAKEN2 classified reads produce relatively more high and medium quality MAGs. The pipeline is open source and available on GitHub (https://github.com/aidanfoo96/MINUUR) with an accompanying JupyterBooks page (https://aidanfoo96.github.io/MINUUR/).

## Materials and Methods

### Specifications

To undertake this analysis, we developed a metagenomics workflow called MINUUR (**M**icrobial **IN**sights Using Unmapped Reads), using the workflow manager Snakemake (40) (Figure 1). This pipeline was developed to facilitate the following analysis and future studies aiming to characterise non-reference reads in mosquitoes or other organisms. A breakdown of the pipeline that produced each result of this study is provided in the following section, with details of how each step was configured in the final section.

**Figure 1:**
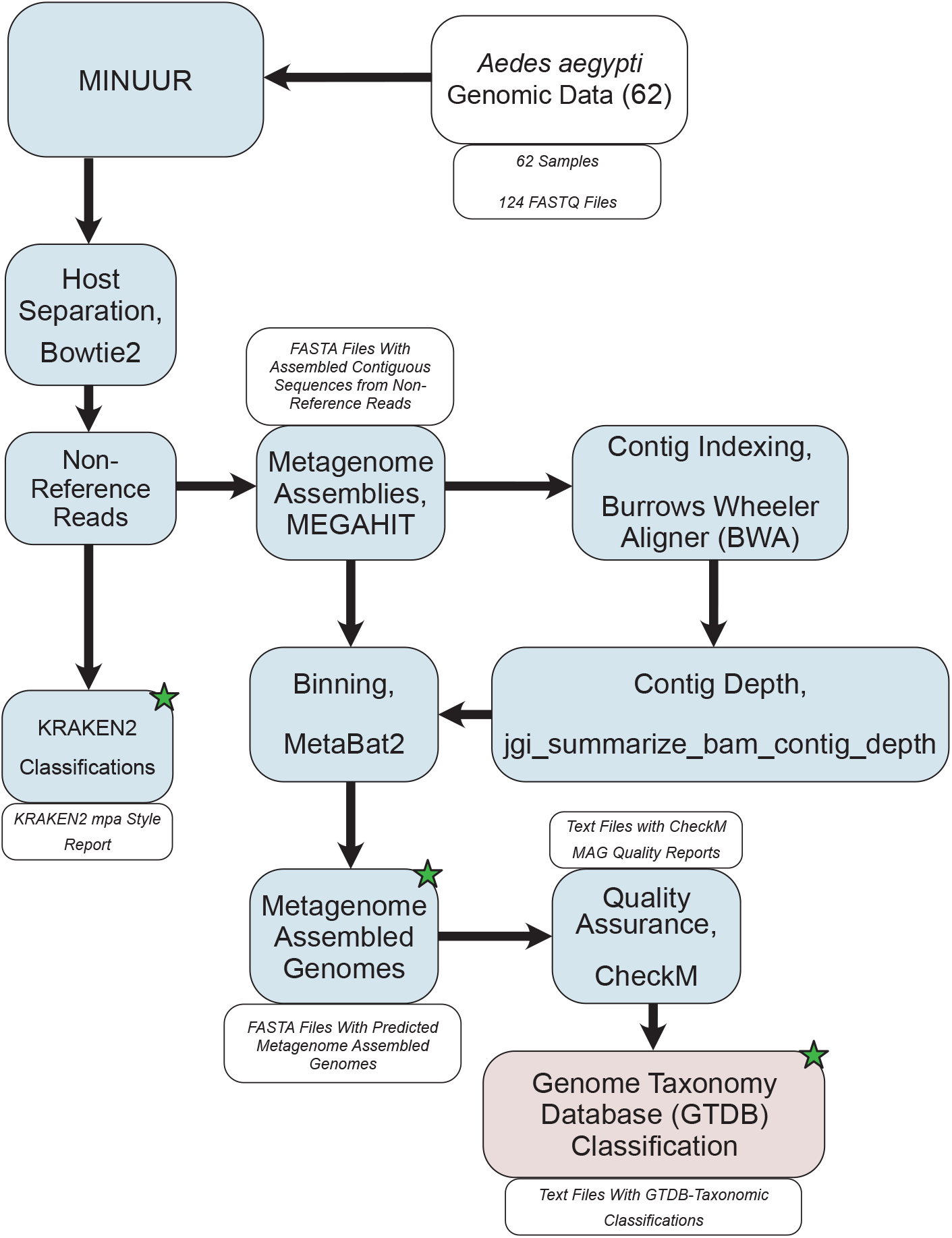
Study Workflow. Workflow describing the main steps of characterising non-reference reads from *Aedes aegypti* whole genome sequencing data. 62 samples (124 FASTQ files) were input to our metagenomics workflow MINUUR (**M**icrobial **IN**sight **U**sing **U**nmapped **R**eads). Reads were mapped to the *Aedes aegypti* reference genome (AaegL5.3) using Bowtie2 (v2.4.4), and unmapped (=non-reference) reads extracted, parsed for taxonomic classification with KRAKEN2 (v2.1.2) and metagenome assembly with MEGAHIT (v1.2.9). The resulting contigs are indexed with burrows wheeler aligner (BWA) (v0.7.17) and used to produce a depth file generated using the jgi_summarize_bam_contig_depth script from MetaBAT2 (v2.12.1). Bins are produced using the assembled contigs and depth file with MetaBAT2. The resulting metagenome assembled genomes (MAGs) are quality checked using CheckM (v1.1.3). High and medium quality MAGs, based on definitions set by the genome standards consortium (65), are taxonomically classified using GTDB-Tk (v1.5.0). Blue boxes denote the steps within MINUUR, red boxes denote steps outside of MINUUR. Green starts denote key outputs of the analyses.

### Database Setup

MINUUR requires several databases to perform the analysis. This includes a high quality bowtie2-indexed reference genome (41) to separate host and non-host reads, and a KRAKEN2 (42), BRACKEN (43) and MetaPhlAn3 (44) database for taxonomic read classifications. All databases are available in their respective GitHub repositories. The databases used in this study are the default MetaPhlAn3 marker gene database, KRAKEN2 and BRAKEN indexes from the Ben Langmead repository located here: https://benlangmead.github.io/aws-indexes/k2. For our study, we downloaded and compiled these default databases on the 08/09/2022.

### Data Preparation

MINUUR accepts either BAM or paired FASTQ inputs. For FASTQ inputs, MINUUR performs quality control (QC) using FASTQC (v0.11.9) (45), providing a QC report per sample. MINUUR does not use the FASTQC report in subsequent steps, but only as a quality assurance metric for the user and to estimate if read trimming is required. Read trimming can be performed within MINUUR using Cutadapt (v1.5) (46) with user defined parameters for minimum read length, base quality and adapter content (default: minimum base length = 50, average base quality = 30). To separate host and non-host, reads are aligned to a user defined indexed reference genome (the relevant host genome) using Bowtie2 (v2.4.4) (41). Alignment sensitivity and type can be adjusted within the pipeline at the user’s discretion. A high quality, chromosome level assembled, reference genome is recommended if available. In situations where this is not possible, users should be aware that read alignment will likely result in mismatches between the reference and target sequence and produce alignments with poor coverage (47). As a result, non-reference reads used in subsequent steps are likely to contain a substantial number of host data. In this instance, we included a feature within MINUUR to extract KRAKEN2 assigned reads pertaining to known microbes and potentially improve metagenome assemblies. Non-reference reads within the coordinate sorted binary alignment (BAM) file are extracted using samtools using the command “ samtools -view -f 4” (v.1.14) (48) and converted to FASTQ format using bedtools (v2.3.0) (49) (“ –bamToFastq”). As users might already have their data in BAM format mapped against the host reference (e.g. in large-scale sequencing projects like the Ag1000G), we also included the option of a BAM input. Here, any non-reference reads within the BAM file will be extracted, converted to FASTQ and used in downstream steps.

For this study, we used five published genome datasets of *Aedes aegypti* publicly available on the European Nucleotide Archive (35,37,50–53). We selected the data sets to cover a range of sequencing depths, DNA extraction method and sequencing platform (Figure 2C). All raw FASTQ files of published sequencing data were retrieved from the European Nucleotide Archive (ENA) under the project accession numbers PRJEB33044 (37), PRJNA255893 (50), PRJNA385349 (51), PRJNA718905 (35), PRJNA776956 (52) and PRJNA992905 (53).

**Figure 2:**
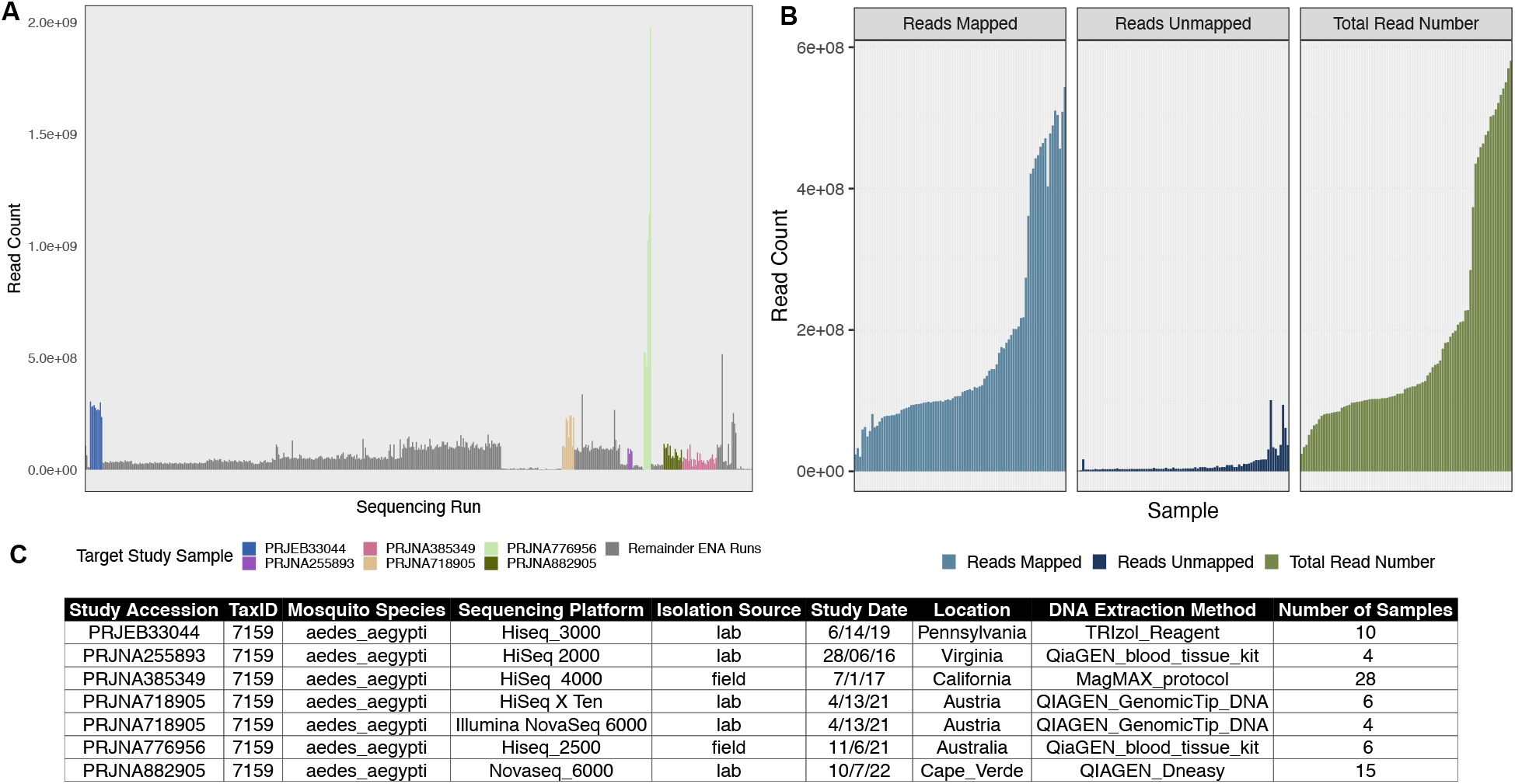
Sequencing Project Selection and Alignment. **A**. Bar chart depicting the total number of whole genome sequencing projects referred to as “ *Aedes aegypti”* on the European Nucleotide Archive. Colour bars show the target sequencing samples we used in our study, colour coded in the legend. **B**. Facetted bar charts showing the total number of reads (right, green), aligned reads (left, blue) and non-reference reads (middle, dark blue) of our selected samples. Mapped and non-reference reads result from the alignment to the *AaegL5*.*3* reference genome using Bowtie2 (v2.4.4) (41) within our metagenomics workflow MINUUR. **C**. Table showing (left to right) the original sequencing project, taxonomic id, mosquito species, sequencing platform, isolation source, study date, location of the sequencing project, DNA extraction method and number of samples.

### Read Classification

MINUUR uses two read classification approaches to infer taxonomy. KRAKEN2 (v2.1.2) (42), which uses a k-mer based approach to map read fragments of k-length against a taxonomic genome library of k-mer sequences, and MetaPhlAn3 (v3.0.13) (44) to align reads against a library of marker genes using Bowtie2 (41). MINUUR also provides the option to use KRAKEN2 classified reads, parsed from KrakenTools (v1.2), to select a specific set of reads (for example bacterial) for metagenome assembly. To estimate the relative taxonomic abundance from KRAKEN2 classifications, MINUUR will parse KRAKEN2 read classifications to BRACKEN (v2.6.2) (43) which uses a Bayesian probability approach to redistribute reads assigned at higher taxonomic levels to lower (species) taxonomic levels.

MINUUR outputs classified and unclassified reads to paired FASTQ files and generates BRACKEN estimated taxonomic abundance profiles for further analysis. An additional feature we included within MINUUR is the option to extract a specific taxon or group of taxa from KRAKEN2, using the program KrakenTools (42). This option is useful in situations where a specific group of taxa are of interest or to exclude groups of taxa such as viral or archaeal reads. Alternatively, if alignment to a reference is poor, this option can be used to remove host reads that did not map to its reference.

### Metagenome Assembly, Binning and Quality Assurance

MINUUR uses the non-reference reads to perform *de novo* metagenome assemblies (the same reads used for KRAKEN2 and MetaPhlAn3 taxonomic classifications). Reads are parsed to MEGAHIT (v1.2.9) (54), a rapid and memory efficient metagenome assembler, for *de novo* metagenome assembly. Assembled contigs are quality checked using QUAST (v5.0.2) (55) to assess contig N50 and L50 scores. The resultant contigs, which are fasta files with sequences pertaining to genomic regions of a microbe, need to be placed within defined taxonomic groups - referred to as a bin. For this, contigs are indexed using the Burrows Wheeler Aligner (BWA) (v0.7.17) (56), and the original non-reference reads are aligned to the indexed contigs using “ – bwa-mem”. The subsequent coordinate sorted BAM file is parsed to the “ jgi_summarize_bam_contig_depth” script from MetaBAT2 (v2.12.1) (57) to produce a depth file of contig coverage. The depth file and assembled contigs are input to the metagenome binner MetaBAT2 (v2.12.1) (57), to group contigs within defined genomic bins. Each bin is a predicted metagenome assembled genome (MAG). CheckM (v1.1.3) (58) is then used for quality assurance of each bin by identifying single copy core genes. Specifically, bin contamination is assessed by looking for one single copy core gene within each bin, and completeness by calculating a required set of single copy core genes.

### Pipeline Configuration

All sample names were listed in the samples.tsv file of the configuration directory and paths to the FASTQ directories given in the samples_table.tsv file. To implement the pipeline, the configuration file was set to the following parameters: FASTQ = True, QC = True, CutadaptParams = “ –minimum-length 50 -q 30”, RemoveHostFromFastqGz = True, AlignmentSensitivity = “ –sensitive-local”, ProcessBam = True, From-Fastq = True, KrakenClassification = True, ConfidenceScore = 0, KrakenSummaries = True, GenusReadThreshold = 1000, SpeciesReadThreshold = 30000, MetagenomeAssm = True, MetagenomeBinning = True, MinimumContigLength = 1500, CheckmBinQA = True. All databases were installed from their respective repositories on 08/09/22. The pipeline was run on an Ubuntu Linux system with 660gb of available memory and 128 CPUs. For our analysis, with the above settings and 10 cores available, runtime was two weeks; the maximum Resident Set Size (RSS) of an individual sample during this run was 9771 RSS (occurring during metagenome assembly); and total storage used (including temporary files) was 5.18Tb (terabytes) across all samples used in this study.

### Taxonomic Classification of MAGs with GTDB-Tk

Separate from MINUUR, all bins produced from MetaBAT2 were taxonomically classified with GTDB-Tk (59) (v2.1.1) using “ –classify-wf” against the Genome Taxonomy Database (GTDB) (release 07-R207 08/04/22, downloaded on 08/09/22). GTDB-Tk assigns genes to MAGs using Prodigal (v2.6.3) (60) and ranks the taxonomic domain of each MAG using a database of 120 bacteria and 122 archaea marker genes (61) using HMMER3 (62). With this information, MAGs are then placed into domain specific reference trees with pplacer (v1.1) (63). Taxonomic classifications with GTDB-Tk are based on placement within the GTDB reference tree, relative evolutionary divergence, and average nucleotide identity (ANI) scores with its closest reference genome. The relative evolutionary divergence score is used to refine ambiguous taxonomic rank assignments and ANI scores are used to define species classifications (59).

## Results

### Extraction of Non-Reference Reads Post-Alignment to *Aedes aegypti*

In total, we retrieved 62 samples (124 FASTQ files) across six sequencing experiments (Figure 2A) and parsed them through MINUUR. After alignment to the *Aedes aegypti* reference genome (AaegL5.3) (64) with Bowtie2, the proportion of mapped and non-reference reads were calculated. Of our initial 62 samples, on average, 176,310,722 reads aligned to the AaegL5.3 reference genome, ranging between 20,655,064 and 543,378,958 reads (Figure 2B). To estimate the number of reads associated to the microbiome among the non-reference reads, the overall number of KRAKEN2 classifications were counted. The average proportion of KRAKEN2 classified reads from all non-reference reads was 17.9%. The number and proportion of KRAKEN2 classifications per sample is given in Supplementary Table 1 within the extended data.

### Taxonomic Classifications Produced from Non-Reference Reads of *Aedes aegypti*

The KRAKEN2 classifications of non-reference reads were counted from each sequencing project. Four out of the six sequencing projects showed a much lower number of classifications compared to sequencing projects PRJNA776596 and PRJEB33044 (Figure 3A). PRJEB33044 gave the highest average proportion of taxonomic classification (81.3%), while PRJNA255893 was the lowest (0.893%).

**Figure 3:**
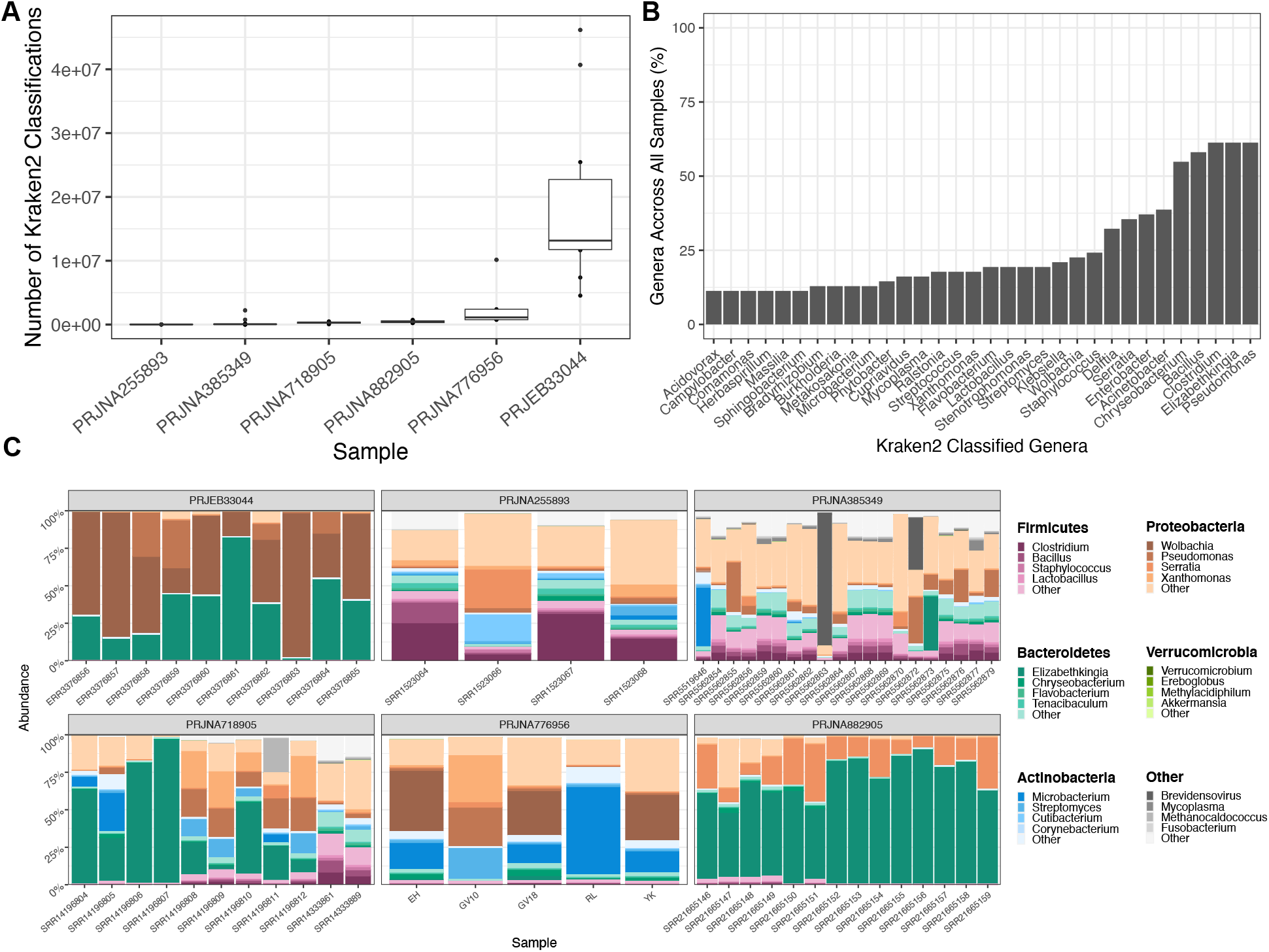
KRAKEN2 (v1.2) Genera Classifications of Non-Reference Reads After Alignment to the *AaegL5*.*3* Reference Genome. **A**. Box plot depicting the number of KRAKEN2 (42) classifications from non-reference reads across 6 publicly available *Aedes aegypti* sequencing projects. Each dot represents one sample within the sequencing project. **B**. Bar chart showing the proportion of genera summed across all samples. **C**. Facetted heatmaps showing phylum and genus level KRAKEN2 assignments of taxa identified from non-reference reads. Colour of each phylum is denoted in the right-hand legend panel, with the most abundant genera depicted by a colour gradient corresponding to their respective phylum. The graph was generated using the microshades R package (78).

Multiple phyla were identified across sequencing projects. *Bacteroidetes, Proteobacteria, Firmicutes and Actinobacteria* were the most common phylum present across all samples, with varied relative abundance between projects (Figure 3C). For example, *Bacteroidetes* and *Proteobacteria* were dominant in PRJEB33044 and PRJNA882905, *Firmicutes* and *Proteobacteria* in PRJNA255893, and *Bacteroidetes, Actinobacteria* and *Proteobacteria* in PRJNA385349 and PRJNA718905. At the genera level, there are several dominant members across samples, including *Wolbachia, Pseudomonas, Serratia* and *Elizabethkingia*. Generally, however, there is considerable variation at the genus level within and between sequencing experiments (Figure 3C). We summarised all KRAKEN2 classifications across 62 samples. The most common genera, identified in >50% of all samples, were *Pseudomonas, Elizabethkingia, Clostridium, Bacillus* and *Chryseobacterium*. Genera identified in 25-50% of samples were *Acinetobacter, Enterobacter, Serratia* and *Delftia* (Figure 3B). All KRAKEN2 taxonomic classifications are given in Supplementary Table 2.

### Metagenome Assembly and Binning

We used MINUUR to further parse non-reference reads through assembly, binning and quality assurance steps, with the aim to recover MAGs associated to the *Aedes aegypti* microbiome. Assembly was conducted with MEGAHIT and binning with MetaBat2. We used CheckM to assess MAG completeness and contamination through the presence and copy number of single copy core genes, and recovered 105 MAGs (Figure 4A). Using the standards of MAG quality set by the genome standards consortium (GSC) (65), 42 MAGs met the criteria of high quality with completeness >90% and contamination <5%; and 20 MAGs classify as medium quality with completeness >50% and contamination <5%. Overall, 62 high and medium quality MAGs were recovered. The remaining MAGs were low-quality (<50% complete and <10% contamination) or contaminated >10%, and therefore excluded from MAG classification. Of the high and medium-quality MAGs, the mean N50 (the minimum contig length of an assembled contig that covers 50% of the genome) was 177.2 kilobase pairs (KBP), ranging between 4.72KBP and 471.6KBP (Figure 4B). The average genome size was 3.36 megabase pairs (MBP), ranging between 1.07MBP and 6.19MBP (Figure 4C), while low-quality MAGs showed a wider range of genome sizes between 0.24MBP and 106MBP (Supplementary Figure 1).

**Figure 4:**
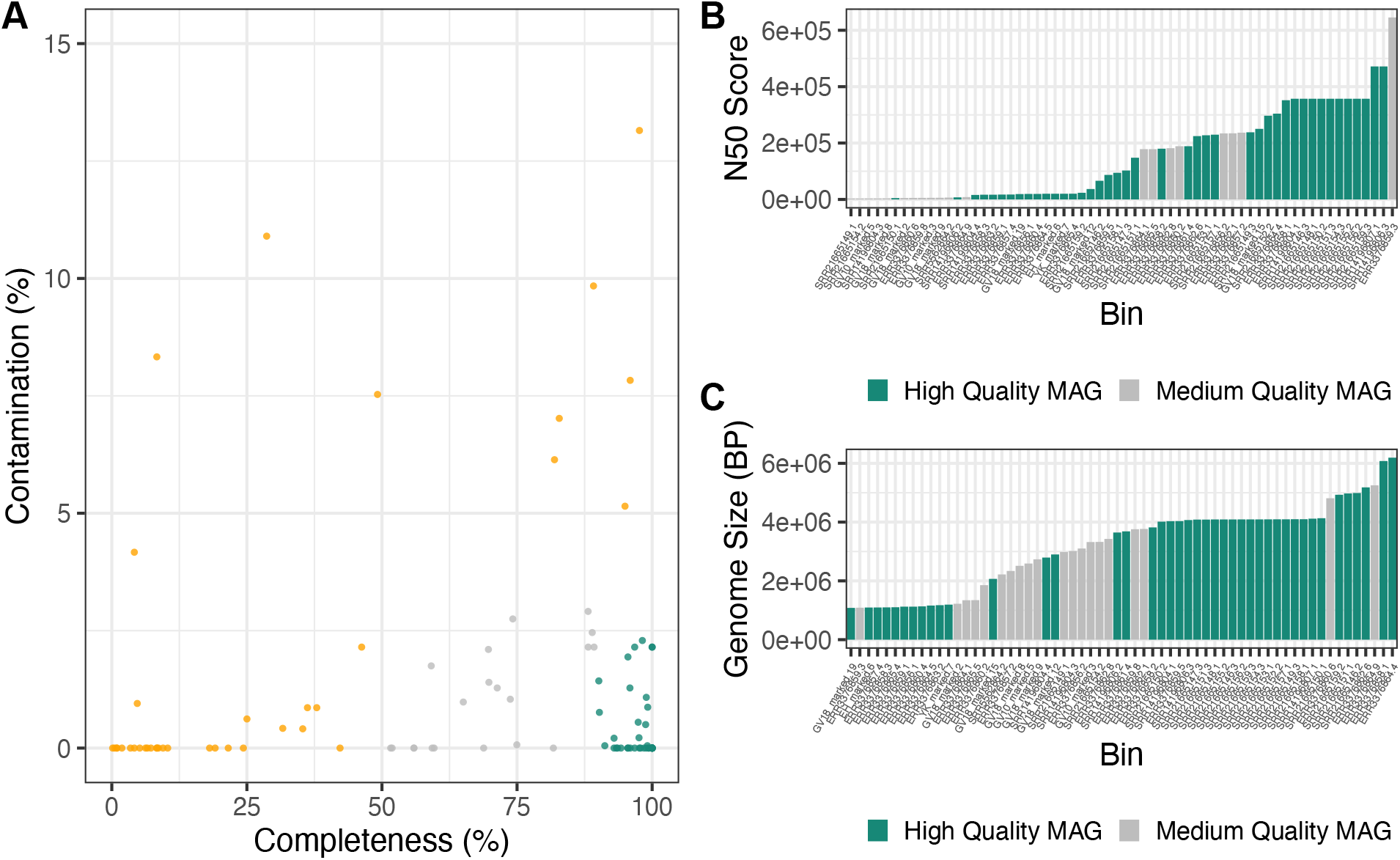
Recovered *Aedes aegypti* Associated Bacterial Metagenome Assembled Genomes (MAGs). **A**. MAGs assembled from non-reference reads using MEGAHIT (v1.2.9) and binned with MetaBAT2 (v2.12.1). Colours denote MAG genome standards consortium (GSC) high, medium and low MAG quality; green = high-quality MAGs (>90% completeness, <5% contamination), grey = medium-quality MAGs (>50% completeness, <5% contamination) and orange = low-quality MAGs. **B**. Bar graph depicting N50 score of medium and high-quality MAGs. Green = high-quality MAG and grey = medium-quality MAGs. **C**. Bar graph depicting genome size in base pairs of medium and high-quality MAGs. Green = high-quality MAG, grey = medium-quality MAG.

### Relationship Between KRAKEN2 Taxonomic Classifications and MAG Recovery

The suitability of a sample for MAG recovery would be of interest to estimate in advance, and we investigated if there was a correlation between the proportion of KRAKEN2 classifications from non-reference reads and MAG recovery. In one project, PRJNA255893, we recovered no high-quality MAGs, (Figure 5A) and the reads used for assembly contained no taxa exceeding 100,000 KRAKEN2 assigned reads (Figure 5B). In contrast, PRJNA33044, PRJNA776596 and PRJNA882905 allowed retrieval of more high and medium quality MAGs, and within these projects, multiple samples contained >100,000 reads assigned to a taxon (Figure 5A, B). As such, MAG recovery is likely linked to a sufficient number of classified reads to a taxon (Figure 5A, Figure 5B), rather than overall number of KRAKEN2 assigned reads within a sample (Supplementary Figure 2). In accordance with this, the total number of classified taxa totaling or greater than 100,000 KRAKEN2 assigned reads for PRJEB33044 is 33 taxa (summed across all samples) and 22 high and medium quality MAGs were recovered from this sample. Similarly, PRJNA776956 shows 19 taxa with associated reads totaling or greater than 100,000 KRAKEN2 assigned reads, resulting in 16 high and medium-quality MAGs. Furthermore, taxonomic classifications with a high number of KRAKEN2 assigned reads (>100,000) are akin to the taxonomic classifications of medium and high-quality MAGs (Supplementary Figure 3). There are, however, exceptions to these patterns; PRJNA385349 contained five taxa with over 100,000 KRAKEN2 assigned reads, yet no high-quality MAGs could be recovered from this project. As such, applicable for future application of this approach, the number of KRAKEN2 classifications assigned to a specific taxon is one factor that can help estimate high-quality MAG recovery.

**Figure 5:**
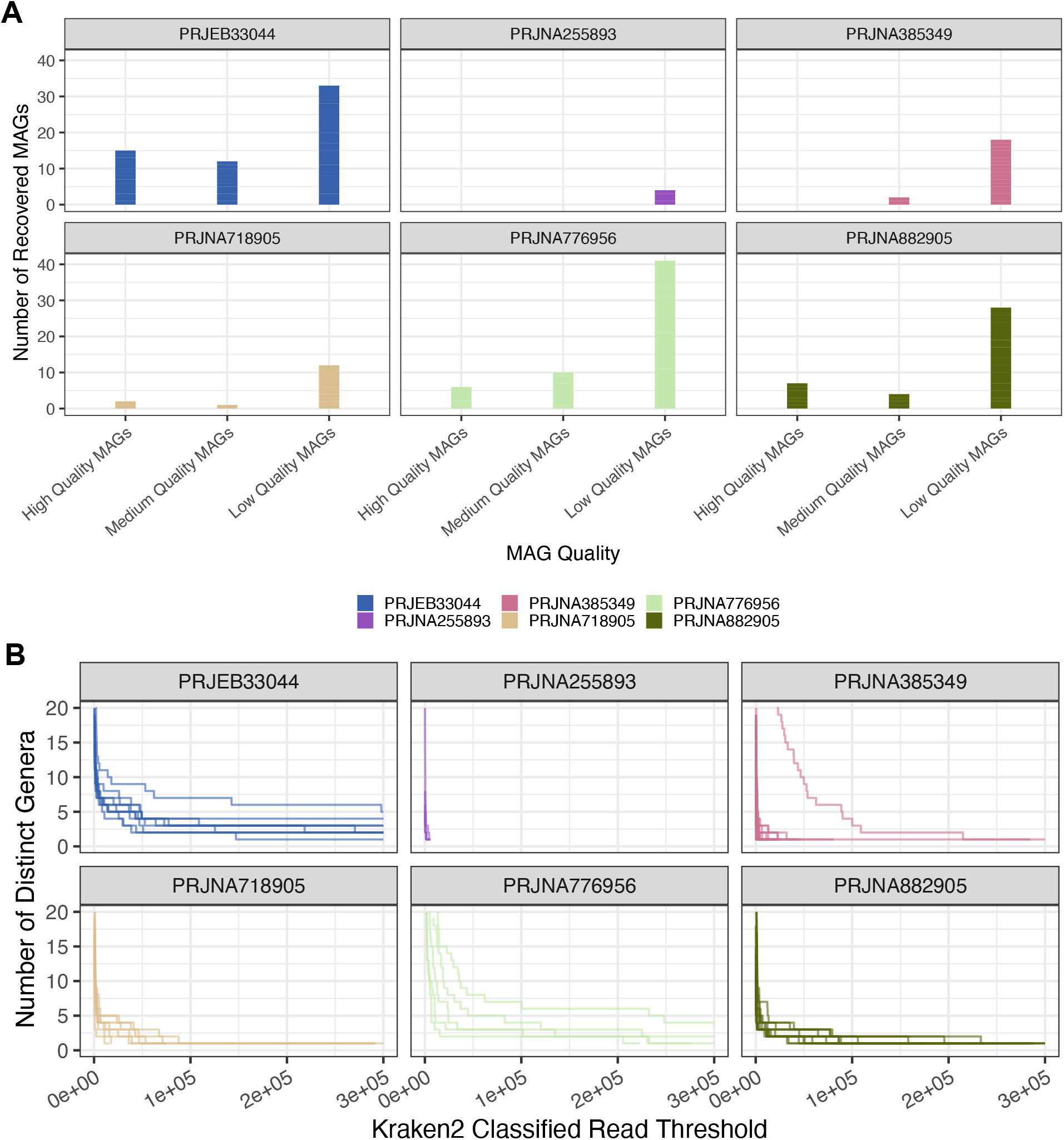
Relationship Between KRAKEN2 classifications and MAG Recovery. **A**. Bar chart depicting the recovery of high, medium and low-quality MAGs from *Aedes aegypti* non-reference reads across each sequencing project. Y-axis = number of recovered MAGs, X-axis = MAG quality ranked using CheckM completeness and contamination. **B**. KRAKEN2 classifications depicted through increased stringency of a KRAKEN2 classification threshold. Each line represents a sample within the sequencing project, Y-axis shows the number of distinct KRAKEN2 classified genera and X-axis represents the number of KRAKEN2 classifications. Each graph represents the filtering of samples towards taxa with a high number of assigned reads.

### Taxonomic Classification of MAGs with GTDB-Tk

Following MAG recovery using MINUUR, we used the taxonomic classifier GTBD-Tk to classify high and medium quality MAGs against the Genome Taxonomy Database (GTDB). We compared the genome size of each MAG to its closest reference genome in the GTDB (Figure 6A, Figure 6B). Of the high-quality MAGs, these were larger than their reference genome by mean = 183kb, whereas medium quality MAGs deviated from their reference genomes by 1768kb (Figure 6A). Congruent with pairwise size differences between MAG and reference genome, we found the overall distribution of high-quality MAGs compared to their reference genome size to be similar, but significantly different between medium-quality MAGs (Figure 6B). Of these, 48 MAGs were classified to the species level with a mean FastANI score of 98.4%, ranging between 95.6% and 100%. No MAGs were identified <95% ANI to a known species, indicating no undescribed *Aedes aegypti* associated bacterial species were present within these MAGs (Figure 6C).

**Figure 6:**
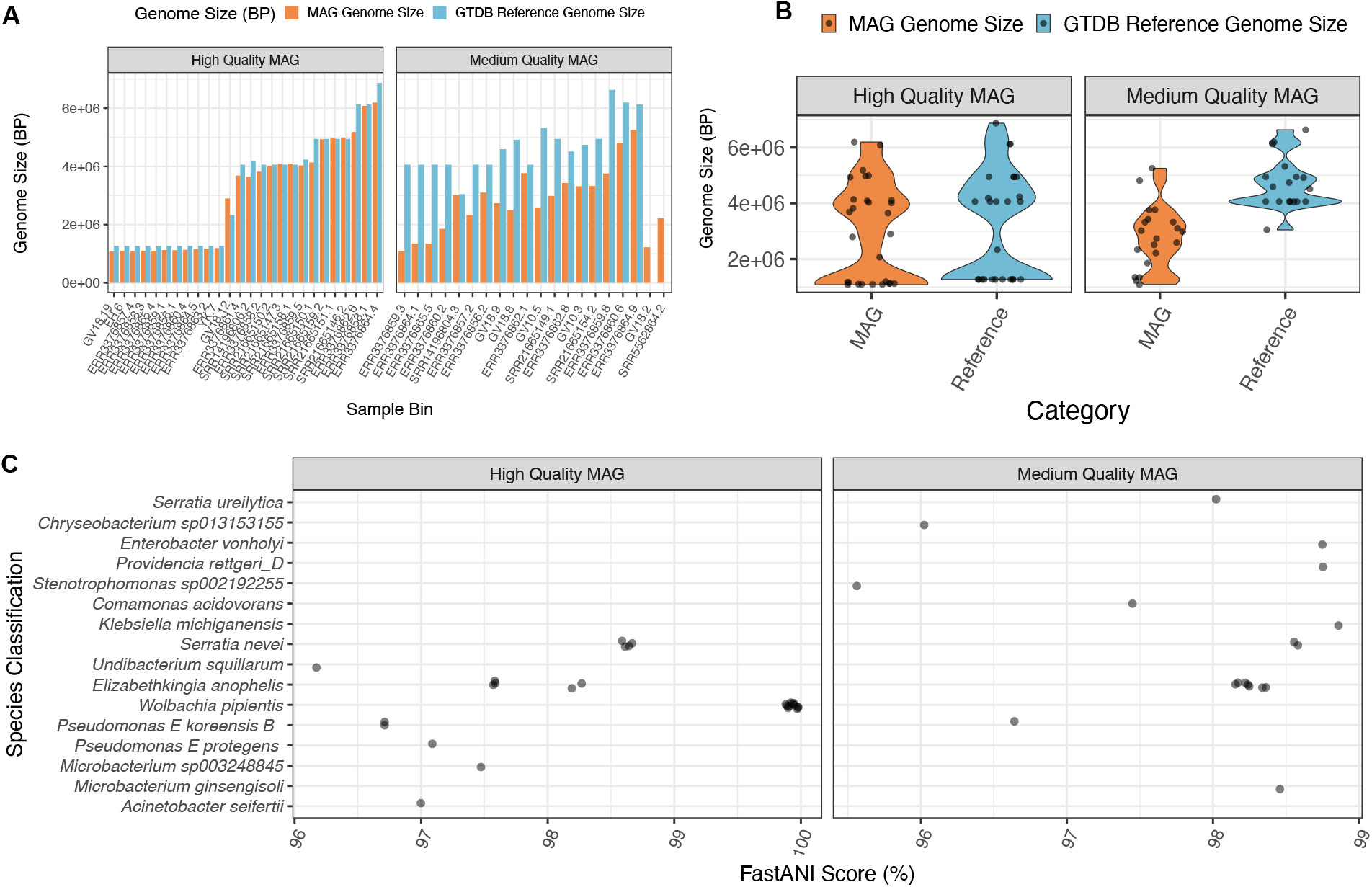
*Aedes aegypti* Associated Bacterial MAG Classifications with GTDB-Tk (v2.1.1). **A**. Genome size (base pairs) of *Aedes aegypti* associated MAGs (orange), split between high-quality (CheckM completeness > 90%, contamination <5%) and medium-quality (CheckM completeness >50%, contamination <5%) rankings, compared to their closest GTDB-Tk reference genome (blue). Each bar denotes the origin sample and the bin number (sample.bin). **B**. Violin plot showing the distribution of *Aedes aegypti* MAG genome size (orange) compared to their GTDB-Tk classified reference genome. Each point denotes a MAG and graphs are facetted between high and medium quality MAGs (see above). **C**. Taxonomic classifications from the Genome Taxonomy Database (GTDB) of MAGs obtained from non-reference *Aedes aegypti* reads. X-axis = FastANI (Average Nucleotide Identity) (%) score to the closest related reference genome in the GTDB, Y-axis = GTDB-Tk species classification.

## Discussion

Metagenomic datasets of mosquito microbiomes are so far limited in number (66). In this study, we developed a metagenomics like workflow called MINUUR to facilitate recovery and use of host-associated bacterial sequences using non-reference reads from existing host WGS projects. We demonstrate that MINUUR can be used to recover genus level taxonomic classifications and draft high and medium quality MAGs from host WGS projects. The recovery of metagenomic information, such as MAGs, are applied in large scale metagenomic studies from chickens (67), humans (68) to cows (69–71), with these studies yielding between 400 to 92,000 MAGs per study. We apply a similar approach with non-reference *Aedes aegypti* sequencing reads across a range of different studies and can demonstrate that using MINUUR expands the genomic representation of known mosquito-associated bacterial symbionts. Overall, these provide a valuable resource for researchers in the field and can be used in further work such as facilitating biosynthetic gene cluster discovery (71) or to identify genetic targets in symbiont pathogen blocking approaches (2).

The data retrieved in this study agrees with published insights; the phylum level classifications are consistent with findings from other mosquito microbiome studies (8,9,11,18,72,73), showing that *Proteobacteria, Bacteroidetes* and *Firmicutes* are dominant phyla of the mosquito microbiome; and our taxonomic classifications highlight the inherent variability of the *Aedes aegypti* microbiome (18,24,74). These findings give us confidence that taxonomic classifications with KRAKEN2, within the pipeline, can accurately predict the presence of microbes associated to the *Aedes aegypti* microbiome from non-reference sequences. At the genus level, we also find consistent observations of taxa previously identified in other studies (16,19,23,75,76). The KRAKEN2 classifications show within the two most common phyla, *Proteobacteria* and *Bacteroidetes*; *Elizabethkingia, Pseudomonas* and *Serratia* are the most common. All three of these symbionts are documented to play key roles in either blood digestion (19,77), iron-acquisition (75) and microbial interactions (16). Notably *Elizabethkingia* has previously been implicated in responses to iron fluxes in *Anopheles gambiae* (75), and blood meals in *Aedes albopictus* (25) and *Aedes aegypti* (24). Similarly, *Pseudomonas* has shown to interact with *Elizabethkingia*, triggering the expression of *hemS* to break down heme into biliverdin catabolites (17). It is encouraging to note the presence of these two bacteria within our taxonomic classifications. We recovered high and medium quality MAGs associated to these taxa (*Serratia, Elizabethkingia* and *Pseudomonas*), which should allow further interrogation of genes associated to these biological processes. To note, the presence of *Wolbachia* in the projects we have analysed is expected since these mosquitoes were transinfected with high titers of this bacterium, further validating the pipeline results (37).

Whilst the nature of the samples can act as confounder given differences in sample handling preparation, we note the relative abundance of the key phyla, and constituent genera, are varied across these projects. Putative biological causes of this variation are likely multifactorial, supported by studies showing environment (9,15), host genetics (23) and competitive mechanisms amongst bacteria (16) to be influential for bacterial colonization in the mosquito.

## Conclusion

In summary, we present a reproducible workflow to analyse host-associated microbial sequence data derived from host WGS experiments, leveraging a vast resource of data for additional insights. Our work focuses on the mosquito microbiome, where future considerations and prospects were recently established by the Mosquito Microbiome Consortium (66). A key point highlighted in this statement was the need for (meta)genomics approaches with solid reproducibility for data analysis within the field. Our pipeline provides a workflow to assess non-host reads from existing mosquito genome sequence data and increases our knowledge of mosquito-associated bacterial genomes. This approach and accompanying workflow will facilitate more analyses of existing WGS data within *Aedes aegypti* and other organisms of interest for the scientific community.

## Data and Software Availability

All original sequencing projects can be accessed under the following accession numbers: PRJEB33044 (https://www.ebi.ac.uk/ena/browser/view/PRJEB33044), PRJNA255893 (https://www.ebi.ac.uk/ena/browser/view/PRJNA255893), PRJNA385349 (https://www.ebi.ac.uk/ena/browser/view/PRJNA385349), PRJNA718905 (https://www.ebi.ac.uk/ena/browser/view/PRJNA718905), PRJNA776956 (https://www.ebi.ac.uk/ena/browser/view/PRJNA776956), PRJNA882905 (https://www.ebi.ac.uk/ena/browser/view/PRJNA882905). The source code for our workflow, MINUUR, is available here: https://github.com/aidanfoo96/MINUUR with an accompanying jupyter books page to run the analysis available here: https://aidanfoo96.github.io/MINUUR/.

## Extended Data

Supplementary Figures and Tables are available in the FigShare repository under the title “ Recovery of Metagenomic Data from the *Aedes aegypti* Microbiome Using a Reproducible Snakemake Pipeline” and can be accessed using the following URL: https://figshare.com/projects/Recovery_of_Metagenomic_Data_from_the_Aedes_aegypti_Microbiome_using_a_Reproducible_Snakemake_Pipeline/158210. This repository contains the following data:

**Supplementary Table 1**: KRAKEN2 Summary Report

**Supplementary Table 2:** KRAKEN2 Per Sample Genera Classifications

**Supplementary Figure 1**: Genome Size Comparison Between High, Medium and Low-Quality MAGs

**Supplementary Figure 2**: Correlation Between Classified Read Number and Number of MAGs

**Supplementary Figure 3**: GTDB Classifications vs KRAKEN2 Assigned Taxa > 100,000 Reads

## Author Contributions

AF, EH, GLH conceived the project. AF and EH designed the methodology. AF performed the analyses and wrote the pipeline. LC provided technical expertise. EH and GLH provided oversight throughout the project. AF and EH drafted the manuscript, and all authors contributed to and approved the final version.

